# Differently sized soluble α-synuclein species from multiple system atrophy and Lewy body disease brains display different seeding propensities

**DOI:** 10.64898/2026.08.03.742519

**Authors:** S. Zampar, Y. Mei, F. Samuel, M. Karadag, I. Martinez-Valbuena, N.R.G. Silver, G. Grimmer, S. Di Gregorio, A. Tandon, G.G. Kovacs, J.C. Watts, M. Ingelsson

**Author notes:** Correspondence: Martin Ingelsson.

## Abstract

Different conformations, or strains, of α-synuclein (α-syn) aggregates are believed to be responsible for the distinct seeding propensities, propagation profiles, and clinical presentations in Lewy body diseases (LBD) and multiple system atrophy (MSA). While biochemical properties and strain differences of insoluble deposits have been extensively characterized, the understanding of what influence soluble α-syn species may have on these processes is limited to a small number of studies focusing on complex mixtures of soluble species or on a single α- synucleinopathy. Given that soluble oligomers are considered highly pathologically relevant, we isolated and characterized the biochemical, seeding, and toxicity properties of size-fractionated soluble α-syn species from MSA and LBD brains, comparing them to species from control brains without known neurological disease (Ctrl). We observed that levels of differently sized oligomers phosphorylated at Ser129, as well as soluble large oligomers (>450 kDa), were increased in LBD compared to both MSA and Ctrl brains. Nevertheless, species derived from MSA brain exhibited seeding activity across the spectrum of α-syn species (oligomers, monomers, and truncated forms) in the seed amplification assay, whereas only oligomeric species (>150 kDa) from LBD cases were seeding-prone. In the HEK293 α-syn (A53T)-YFP biosensor line, as well as in murine primary neurons, only large oligomers (>450 kDa) from MSA cases induced seeding and aggregation of α-syn. Taken together, our study suggests that soluble α-syn species derived from MSA and LBD brains show different biochemical, aggregation and seeding patterns, presumably due to strain variations of the respective oligomers. Our findings provide novel insight into the pathogenesis of different α-synucleinopathies, which may guide us in the development of targeted therapeutics.

## Introduction

Aggregation and accumulation of α-synuclein (α-syn) in the brain are common features of a family of neurodegenerative diseases referred to as α-synucleinopathies, which include Parkinson’s disease (PD), dementia with Lewy bodies (DLB), and multiple system atrophy (MSA). In PD/DLB (Lewy body diseases, LBD), α-syn deposits are primarily found as Lewy bodies/neurites in neuronal cells but are also present in astrocytes and oligodendroglia (1, 2). In MSA, glial cytoplasmic inclusions of α-syn are mainly present in oligodendrocytes but can also be observed in neurons (1).

Over the past decade, structural studies have demonstrated that α-syn aggregates can adopt disease-specific conformations, or “strains” (3–5), and that such structural differences translate into distinct seeding, propagation, and toxicity profiles. By seeding, we refer to the ability of pathogenic protein forms to transfer their conformation to physiological species, ultimately leading to the propagation of brain pathology (6). The strain hypothesis is supported by a growing body of *in vitro* and *in vivo* studies (7, 8). In transgenic mouse brains, inoculation with MSA and LBD brain extracts elicits strain-specific seeding and propagation behaviors (9), which can be retained upon serial passaging in transgenic mice (10).

While insoluble deposits are the neuropathological hallmarks of these diseases, it has been shown that oligomeric α-syn is particularly neurotoxic (11–13) (and reviewed in (14)). Studies examining oligomeric species extracted from human *post mortem* brain tissue, using both in cell and animal models, have provided key insights into their pathogenicity (15). Consistent with this, oligomers generated from recombinant α-syn have been found to efficiently seed α-syn monomers *in vitro* (16, 17), and to drive propagation and toxicity both *ex vivo* (17) and *in vivo* (18).

In addition to full-length α-syn, truncated α-syn species have been implicated in the disease pathogenesis across α-synucleinopathies (19). C-terminal truncations have been shown to increase α-syn aggregation propensity and drive neurodegeneration and motor dysfunction in a mouse model (20), whereas the less abundant N-terminal truncations have been identified as a characteristic post-translational modification (PTM) of astrocytic α-syn deposits in LBD brains (21). Next to truncation, several studies have suggested that phosphorylation of α-syn at Ser129 plays a central role in α-synucleinopathies. While less than 5% of α-syn was found to be phosphorylated under physiological conditions, pSer129-syn accounts for approximately 90% of α-syn in pathological inclusions, both in LBD and MSA (22, 23). Nonetheless, the precise contribution of this PTM to α-syn aggregation, seeding, and toxicity remains incompletely understood (22, 24).

Notably, most studies investigating α-syn seeding and propagation have relied on synthetic or recombinant protein preparations, despite evidence that brain-derived α-syn variants diverge structurally from their recombinant counterparts (5, 25) and display distinct aggregation behaviors in cell models (26). When human brain-derived samples have been used, most studies have employed fibrillar aggregates or preparations likely containing a heterogeneous mix of assemblies that vary in size and solubility, making it challenging to determine the seeding properties for differently sized species. Consequently, although fibril strains are well characterized, it remains unclear whether oligomeric α-syn species also exhibit disease-specific differences and to what extent they contribute to seeding within the diseased brain.

In this study, we aimed to characterize the seeding properties and toxic effects of differently sized soluble α-syn species isolated from *post mortem* brains. In addition to MSA and LBD cases, we included control brains without known neurological disease (Ctrl). By employing size-exclusion chromatography (SEC), we isolated soluble oligomers and truncated α-syn species from PBS homogenates of temporal cortex samples and identified differences between LBD and MSA brain-extracted oligomers, both *in vitro* and *ex vivo*. Collectively, our findings highlight the heterogeneity of soluble brain-derived α-syn and advance our understanding of which species are most pathogenic across the α-synucleinopathies.

## Results

### Phosphorylation at Ser129 and aggregation state differentiate soluble α-syn species from MSA, LBD and Ctrl brains

The crude PBS brain homogenates (BH) from *post mortem* human MSA, LBD and Ctrl cases (demographic information summarized in Table S1) were centrifuged to remove large insoluble aggregates (i.e. α-syn fibrils). The total protein amount of the cleared PBS-soluble homogenates was normalized and subjected to SEC in order to isolate soluble α-syn species based on their size. Twenty-seven fractions were collected for each case, starting 1.5mL before the column void volume (V_0_), which was identified by Dextran Blue injection (Suppl. Fig.1A). Recombinant α-syn was run on the same column to identify the elution time of monomers, which elution peaked in fraction 18 (Suppl. Fig. 1B). Total protein and total α-syn levels were measured in each fraction and were detectable in fractions 4-27. Accordingly, differently sized soluble α-syn oligomers eluted in fractions 4-17, whereas truncated α-syn species eluted in fractions 19-27.

**Fig. 1.**
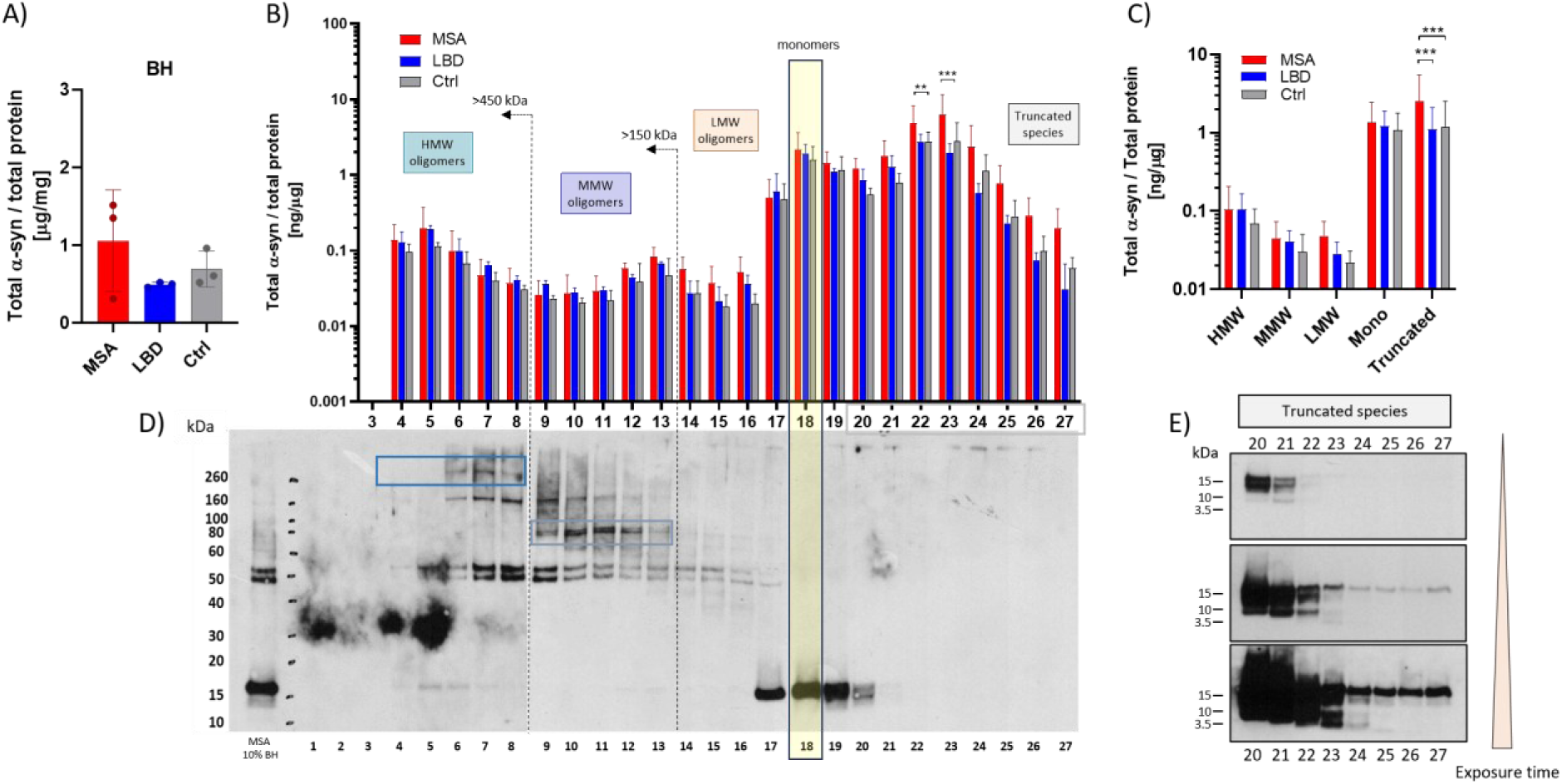
Separation of soluble α-syn species in MSA, LBD, and Ctrl brains using size exclusion chromatography. **A)** ELISA-based quantification of total α-syn in cleared PBS-BH from MSA, LBD and Ctrl brains. **B)** ELISA-based quantification of total α-syn in each SEC fraction, expressed relative to total protein content, compared between MSA, LBD and Ctrl. **C)** Total α-syn levels in relation to total protein levels in grouped SEC fractions corresponding to high molecular weight (HMW, fractions 4-8), mid-sized molecular weight (MMW, fractions 9-13), low molecular weight (LMW, fractions 14-16) oligomers, monomers (Mono, fractions17-19) and truncated species (fractions 20-26) **D)** Representative denaturing western blot for total α-syn across SEC fractions. Immunoblot showing the elution profile of α-syn species, illustrating the distribution of α-syn forms of different sizes across SEC fractions. **E)** Denaturing western blot developed with extended exposure times, revealing lower-molecular-weight truncated α-syn species enriched in specific SEC fractions. Data are given as means ± SD. Two-way ANOVA followed by Bonferroni’s multiple comparison test: *p < 0.05, **p < 0.01, ***p < 0.001.

Total α-syn levels were measured via ELISA in the starting BH and in each SEC fraction. No significant differences were observed between of MSA, LBD, and Ctrl cases (Fig. 1A). Comparable levels of monomeric and oligomeric α-syn were observed among MSA, LBD, and Ctrl brains after fractionation (Fig. 1B,C). In contrast, truncated α-syn species showed increased levels in fractions 22–23 of the MSA group compared with Ctrl. Calibration standards were used to interpolate the molecular weight (Mw) of the proteins eluted from the SEC column (Suppl. Fig.1C-E). As previously reported (27, 28), the separation of α-syn and other intrinsically disordered proteins does not follow the same pattern as the globular standard proteins, and elutes at a fraction with a higher interpolated molecular weight than the expected 14.5 kDa. Thus, the extrapolated Mws for the differently-sized α-syn oligomeric species we report in this study are approximations.

Visualization of α-syn across different SEC fractions under denaturing conditions revealed distinct band patterns that were used to group α-syn species (Fig. 1D). Fractions 4-8 showed a unique band running above the 260 kDa ladder standards, which corresponded to high molecular weight (HMW) oligomers larger than 450 kDa. Fractions 9-13 displayed a distinct band at approximately 80 kDa, which corresponded to mid-sized molecular weight (MMW) species between 450 and 150 kDa. Fractions 14-16 displayed weak α-syn immunoreactivity, which corresponded to low molecular weight oligomers (LMW) species between 150-29 kDa. Fraction 18 contained monomers, in agreement with the elution profile of recombinant α-syn. Finally, fractions 20-27 contained differently-sized truncated α-syn species. The two fractions shouldering the peak elution of monomer in fraction 18 contained a mixture of monomers and LMW oligomers (fraction 17) or truncated species (fraction 19). For fractions 20-25, a separate SEC run with doubled starting protein input and longer exposure times was required to visualize bands by Western blot (WB, Fig. 1E), showing bands at 12 and 10 kDa (fractions 20-22), and below 3.5 kDa (fractions 23-25). For fractions 26 and 27, α-syn immunoreactivity was detected by ELISA, but no truncated α-syn bands were observed by WB under these experimental conditions.

Since total α-syn levels did not differ among the cases, except in a few fractions containing truncated species, we next investigated whether the extent of α-syn phosphorylation or its aggregation state changed between disease and control brains. To measure pSer129 α-syn and aggregated α-syn levels, we adapted an electrochemiluminescence assay on the MSD platform using the pSer129-specific monoclonal antibody (mAb) MJFR-13 and the conformation-specific mAb MJFR-14, binding to aggregated α-syn (Table S2, Suppl. Fig. 2).

No significant differences in pSer129 α-syn levels, normalized to total α-syn, were observed in whole brain homogenates between MSA, DLB and Ctrl cases (Fig. 2A), but analysis of the differently sized soluble α-syn species revealed that oligomeric species of LBD cases are more frequently phosphorylated compared to MSA and Ctrl cases, an increase that reached statistical significance for specific fractions (Fig. 2B). Although truncated and monomeric α-syn species showed low relative phosphorylation, their absolute pSer129 α-syn levels (normalized to total protein) exceeded those observed in oligomeric or monomeric fractions. (Suppl. Fig. 3A). The Ctrl group exhibited the highest overall pSer129 α-syn levels in fractions corresponding to truncated species. For the LBD group, decreased levels of pSer129 α-syn were measured in fractions 22- 24 compared to the Ctrl group. Dot-blot analysis of pSer129 in the different SEC fractions aligned with the MSD ELISA finding, confirming differential phosphorylation in the LBD group with regard to oligomeric and truncated α-syn species (Suppl. Fig. 3B,C).

**Fig. 2.**
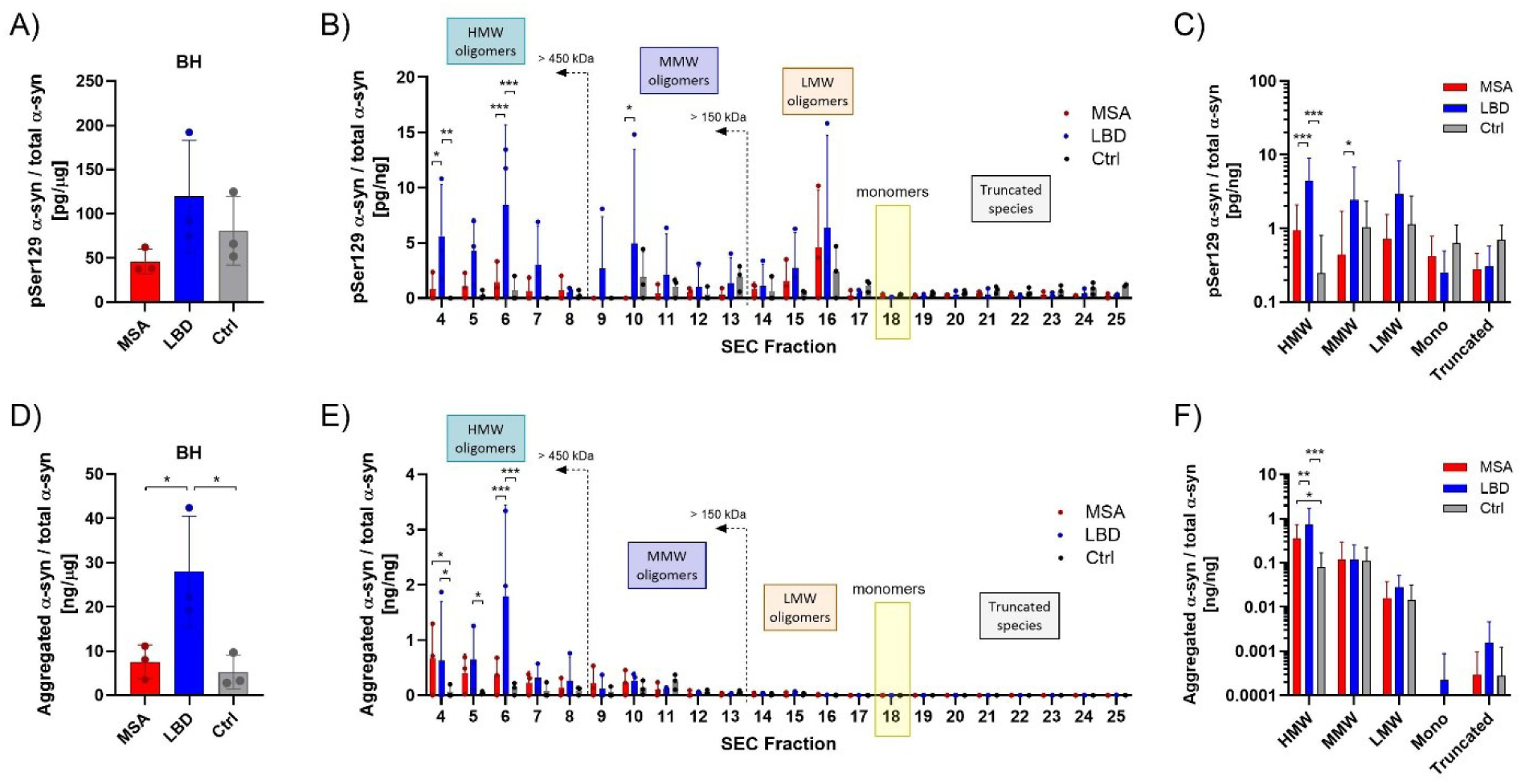
Measurements of pSer129 and aggregated α-syn species in MSA, LBD, and Ctrl brains. **A)** MSD-based quantification of pSer129 α-syn in Ctrl, LBD, and MSA brain homogenates, normalized to total α-syn. **B)** Quantification of pSer129 α-syn in isolated SEC fractions, normalized to total α-syn comparing Ctrl, MSA and LBD cases. **C)** pSer129 α-syn levels in grouped SEC fractions corresponding to high molecular weight (HMW, fractions 4-8), mid-sized molecular weight (MMW, fractions 9-13), low molecular weight (LMW, fractions 14-16) oligomers, monomers (Mono, fractions 17-19) and truncated species (fractions 20-26). **D)** Aggregated α-syn levels in brain homogenates, normalized to total α-syn. **E)** Aggregated α-syn levels across SEC fractions, normalized to total α-syn. **F)** Aggregated α-syn levels in grouped SEC fractions corresponding to high molecular weight (HMW, fractions 4-8), medium molecular weight (MMW, fractions 9-13), low molecular weight (LMW, fractions 14-16) oligomers, monomers (Mono, fractions 17-19) and truncated species (fractions 20-26). Data are given as means ± SD. Two-way ANOVA followed by Bonferroni’s multiple comparison test: *p < 0.05, **p < 0.01, ***p < 0.001.

Measurement of aggregated α-syn levels with the conformation-specific MJFR14 mAb also highlighted differences between the groups. The LBD brain homogenates had significantly higher relative aggregated α-syn levels than MSA and Ctrl brains (Fig. 2D). As expected, when analyzing species of different sizes, HMW oligomer-containing fractions had the highest levels of aggregated α-syn (Fig. 2E,F). Although both MSA and LBD had increased levels of relative aggregated α-syn in HMW fractions compared to Ctrl, LBD exceeded the MSA group (Fig. 2F), with the highest levels observed specifically in fraction 6 (Fig. 2E). Brains derived from Ctrl individuals had overall low α-syn aggregates, even in HMW-oligomer-containing fractions. The reactivity progressively decreased through MMW-oligomers, and signals were below the quantification or detection limit from fraction 16 onwards. The absolute abundance of aggregated HMW α-syn was not statistically different between MSA and LBD fractions, which, however, were increased compared to non-diseased individuals (Suppl. Fig. 3D). Consistent with the MSD assay, dot-blot analysis showed aggregated α-syn immunoreactivity most prominently in fractions 4–8, declining in subsequent fractions and absent, as expected, in the monomer-containing fraction 18. In this semi-quantitative analysis, however, no significant differences in aggregated α-syn were detected among MSA, LBD, and Ctrl groups (Suppl. Fig. 3E,F).

### Brain-derived MSA and LBD soluble oligomers show different seeding profiles *in vitro*

To investigate the seeding propensities of the differently-sized soluble α-syn species, we first performed *in vitro* seed amplification assays (SAA). The fitted curves were used to derive the half-time of aggregation (T50), defined as the time required for the fitted fluorescence signal to reach 50% of its maximal amplitude, and the area under the curve (AUC), calculated by integrating the fitted function over the duration of the assay, which served for kinetic comparison across samples or conditions. Under the chosen assay conditions, SAA using cleared PBS-soluble brain homogenates showed different seeding propensities between MSA, LBD, and Ctrl, as previously reported (29), with lower T50 in the MSA and LBD groups and increased AUC for MSA (Fig. 3A,B). Despite the presence of the aggregation-reducing K23Q mutation in the α-syn substrate (template), we observed frequent spontaneous aggregation of the template during the SAA in buffer-only reactions, with substantial inter- and intra-lot variability (Suppl. Fig. 4A,B). Consistent with the proposed inhibitory role of biological matrix components, we observed that total protein levels in the seed sample affected spontaneous aggregation, with a positive correlation between total protein content and T50 (Suppl. Fig. 4C), i.e., lower amounts of total protein in the seed sample generated faster aggregation kinetics. SAAs are influenced not only by the concentration and conformational properties of pathological α-syn seeds but also by the biological matrix in which they are present. Endogenous proteins, including albumin, lipoproteins, molecular chaperones, and other soluble matrix components, have been shown or proposed to modulate fibril nucleation and elongation, thereby influencing assay kinetics and sensitivity (30–32). Furthermore, control brain homogenates have been found to suppress spontaneous α-syn aggregation compared with unseeded recombinant reactions, suggesting that endogenous matrix proteins inhibit nonspecific nucleation (33). Accordingly, dilution of brain preparations may reduce the concentration of these inhibitory factors, lowering the threshold for spontaneous template aggregation and thereby contributing to altered SAA kinetics. To determine the concentration threshold that would prevent or attenuate spontaneous template aggregation, we examined serial dilutions of brain homogenate from C57BL/6 and triple-syn KO mice in SAA reactions (Suppl. Fig. 4D,E), which revealed that in-assay protein concentrations ≥20 μg/mL prevented nonspecific template aggregation. Because the isolated SEC fraction’s protein content ranged between 10- 100 μg/mL (Suppl. Fig. 5A), and normalization of α-syn levels would lead to ≤10 μg/mL total protein content, we spiked the SAA reactions with triple-syn KO homogenate at a concentration of 40 μg/mL. Without such addition, a strong positive correlation between total protein concentration and T50 was also observed in SAA reactions using the SEC fraction, regardless of the experimental group (Suppl. Fig. 5B,C).

**Fig. 3.**
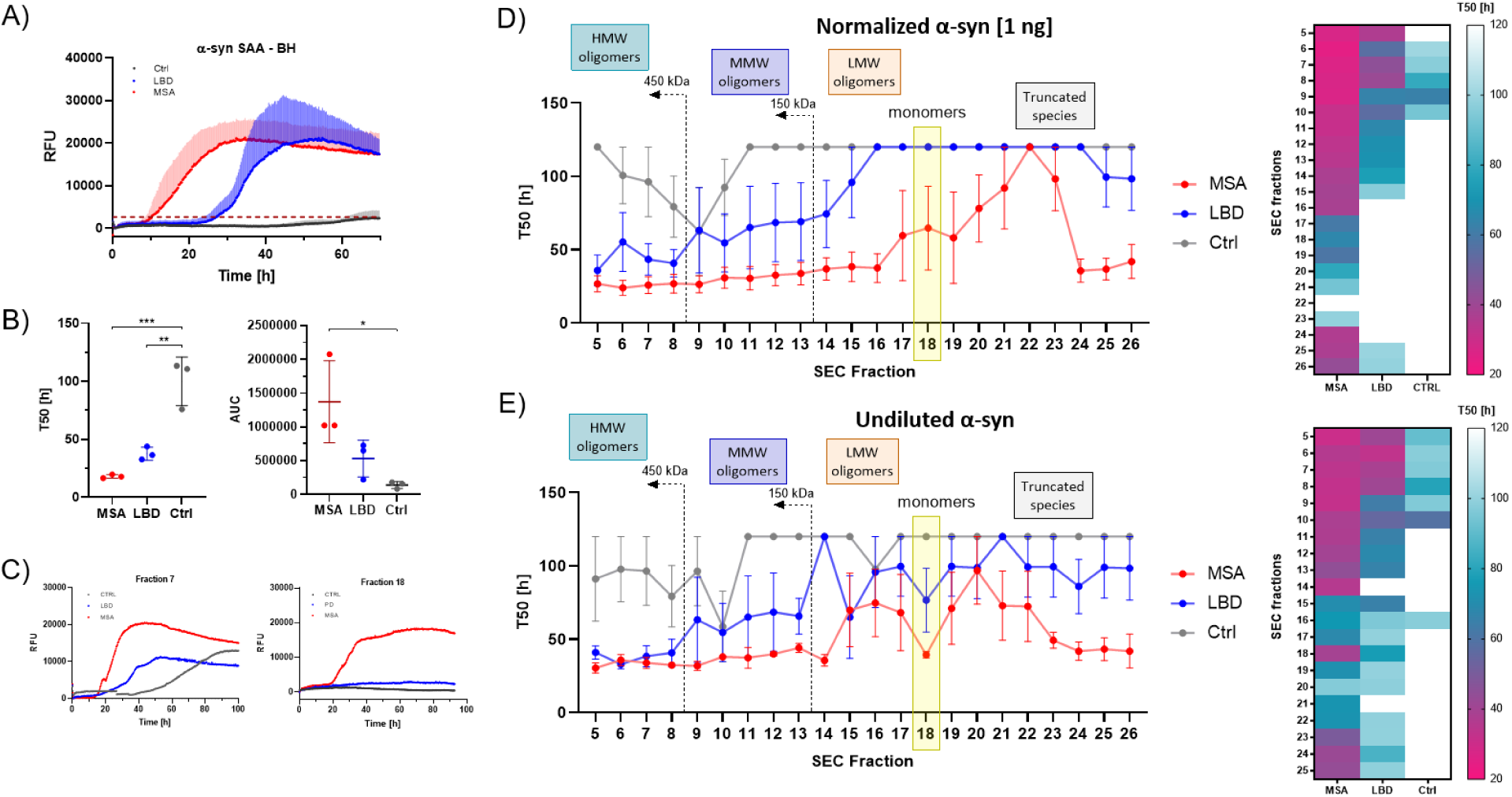
Seed amplification assay reveals distinct seeding profiles of brain-derived soluble α-syn species. **A)** Representative SAA aggregation curves from brain homogenates. Real-time fluorescence traces from MSA, LBD and Ctrl brain homogenates illustrating disease-specific differences in seeding potency and aggregation kinetics. **B)** Half-time (T50) and area under the curve (AUC) from BH-derived SAA reactions. **C)** Representative SAA curves from SEC fraction 7 (HMW oligomers) and fraction 18 (monomers), showing case-specific differences in seeding activity. **D)** T50 across SEC fractions in SAA after normalization to 1 ng α-syn and addition of triple-syn KO BH to normalize total protein content. Heatmap showing fraction-wise T50 values for MSA, LBD, and Ctrl. Lower T50 values indicate faster aggregation and stronger seeding activity. **E)** T50 across SEC fractions in SAA without α-syn normalization and with addition of triple-syn KO BH to normalize total protein content. Heatmap displaying raw (non-normalized) T50 values across fractions for MSA, LBD, and Ctrl. Data are given as means ± SD. One- way ANOVA followed by Bonferroni’s multiple comparison test: *p < 0.05, **p < 0.01, ***p < 0.001.

To assess potential strain differences between groups, the SAA reactions were initially performed with α-syn content normalized to 1 ng per seeding reaction. Despite identical α-syn input, the seeding propensities differed remarkably between diseased and control cases. Fractions derived from MSA brains induced robust seeding across the analyzed α-syn size range, except for some fractions in the truncated species pool (21–23), as indicated by low T50 values (Fig. 3D). In the LBD group, only HMW and MMW oligomers were seeding-positive, but with more variable and higher T50 values than in MSA fractions. The α-syn species isolated from Ctrl cases were largely seeding-incompetent, except for low to moderate seeding reactions occurring in oligomer-containing fractions 6-10. These data highlight strain-dependent differences in seeding behavior at the level of α-syn oligomers. While α-syn normalization allowed such a comparison, the overall amount of α-syn differed across the fractions (Fig. 1A). To assess the seeding propensities of the differently sized α-syn species while maintaining their “original” relative abundance, we next performed SAA on undiluted fractions (Fig. 3E). The overall seeding profile was comparable to the normalized seeding reactions, with MSA-derived α-syn species of different sizes being highly seeding competent, fractions from LBD cases showing decreasing seeding capacity with decreasing sizes (HMW>monomers>truncated species), and only HMW α-syn from Ctrl cases showing moderate seeding activity. No significant correlation was observed between total α-syn levels and T50 in the LBD or Ctrl groups, while a positive correlation was detected in MSA (Suppl. Fig. 5D). This suggests that total α-syn abundance does not predict seeding activity, consistent with the notion that seeding is driven by the presence of aggregation-competent species rather than the overall amount of α-syn. Breaking down the single fractions’ contribution, it became indeed evident that HMW oligomeric fractions, despite having the lowest amount of α-syn, are the most seeding-prone, followed by MMW oligomer-containing fractions, in diseased cases. Monomers and the first-appearing truncated species (fractions 19-22), despite containing around 30 times more α-syn per mg of total protein, contribute little to the seeding behavior of soluble α- syn (Suppl. Fig. 5E).

### HMW oligomers from MSA cause seeding and aggregation in cell models

We next analyzed the seeding behavior of the isolated α-syn species in cellular models. We first employed a well-characterized HEK293 biosensor line expressing A53T-α-synuclein-YFP, previously used for inclusion formation assays (8). When seeding competent species, such as α- syn preformed fibrils (PFF), is applied to the cell line, a fluorescent cytosolic inclusion develops as a consequence of seeded aggregation of A53T-α-syn-YFP. Our assessment of cleared PBS brain homogenates was consistent with previous findings (8), showing that YFP-inclusion-positive cells were observed only in MSA-treated cells. While LBD-treated cells were generally negative, a few thread-like inclusions were occasionally observed (Suppl. Fig. 6A,B). When differently sized soluble α-syn species from MSA brains were applied to the biosensor line, we observed seeding reactions occurring only with HMW oligomers, with the highest percentage of YFP-inclusions after exposure to fractions 6-7 (Fig. 4A,C). Cells treated with LBD brain-derived fractions were largely negative, but for a few thread-like aggregates forming with HMW oligomers in fraction 7. Such inclusions appear to differ in shape from those in MSA-seeded reactions, which formed globular aggregates (Fig. 4B). Soluble α-syn species isolated from Ctrl brains did not lead to any formation of inclusions.

**Fig. 4.**
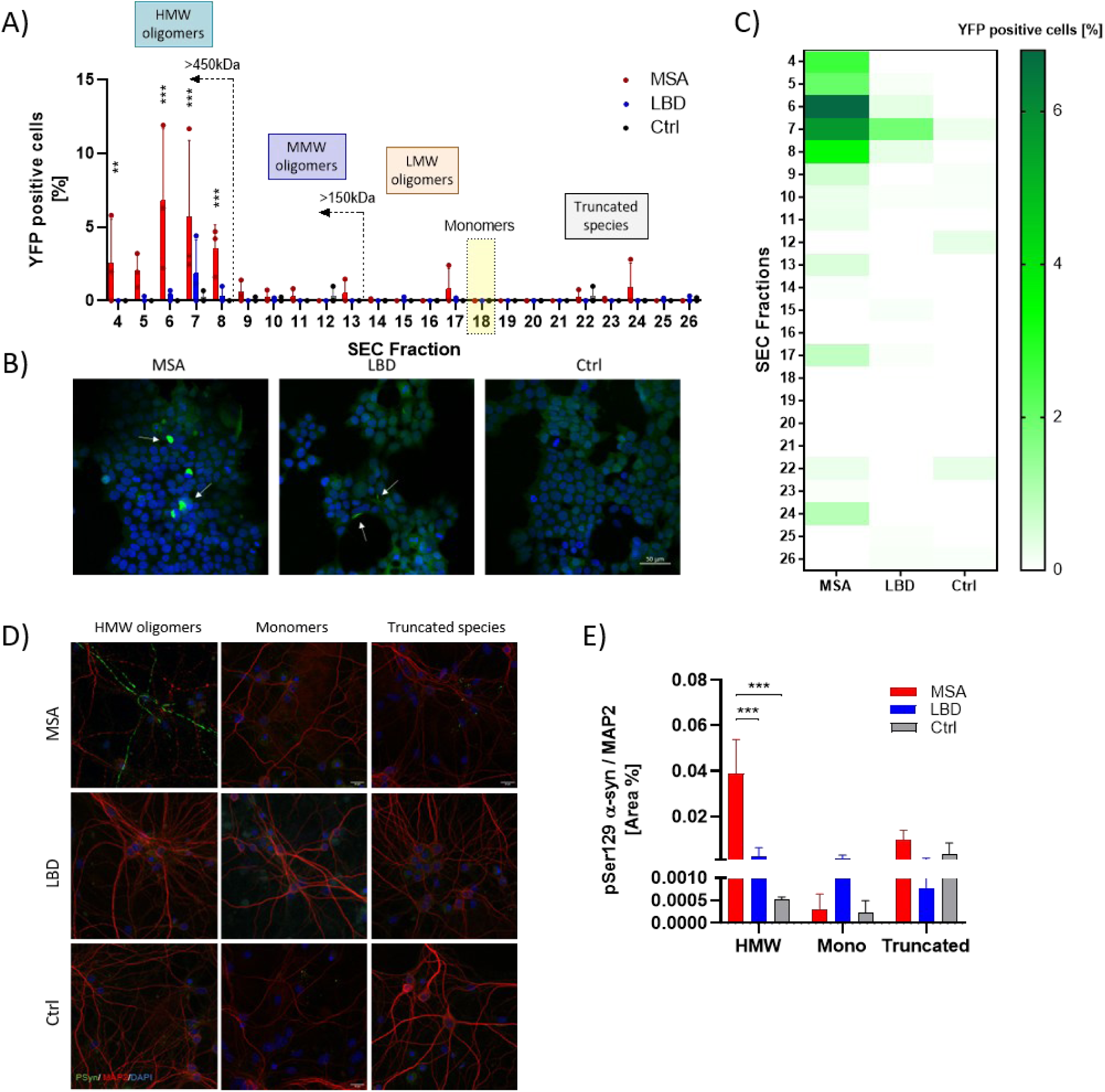
HMW brain-derived α-syn oligomers from MSA seed inclusions in HEK293 biosensor cells and murine primary neurons. **A)** Quantification of seeding activity in HEK293 A53T-α-synuclein-YFP biosensor cells exposed to SEC fractions. Percentage of YFP-positive cells across all fractions from MSA, LBD and Ctrl brains, demonstrating fraction-dependent seeding potency and highlighting enhanced activity in high-molecular-weight fractions for the MSA group. **B)** Representative biosensor images from SEC fraction 7, showing robust intracellular YFP-positive aggregates in cells treated with MSA samples. Scale bar: 50 µm. **C)** Heatmap of seeding activity across SEC profiles, illustrating seeding intensity for MSA, LBD and Ctrl samples. **D)** Representative images of primary neurons treated with pooled SEC fractions - HMW oligomers (4–8), monomers (18), and truncated species (21–24) - showing pSer129-positive inclusions (green) after exposure to HMW species from MSA. Microtubule-associated protein (MAP2, red) marks neuronal morphology; DAPI (blue) counterstains nuclei. Scale bar: 50 µm. **E)** Quantification of pSer129 α- syn signal in primary neurons after SEC-fraction treatment. Data are given as means ± SD. Two-way ANOVA followed by Bonferroni’s multiple comparison test: **p < 0.01, ***p < 0.001.

We next analyzed the seeding capacity of soluble α-syn species in a more physiological model by exposing murine wild-type primary neurons to different SEC fractions. We observed robust pS129-syn immunoreactivity upon application of recombinant α-syn fibrils to neuronal cultures (Suppl Fig. 6C), and exposure to whole PBS-solubile brain homogenates led to the formation of pS129-syn-immunoreactivity only with MSA cases (Suppl. Fig. 6D). We then pooled fractions containing HMW-oligomers (4–8), and truncated species (21–24) and compared their *ex vivo* seeding activity to monomers (fraction 18). Consistent with the finding from the biosensor line, only HMW oligomers from MSA brains seeded and induced the accumulation of pSer129 α-syn, while monomers and truncated species, irrespective of their source, were inert (Fig. 4D,E).

### Toxicity of differently sized α-syn species

To compare the toxicity of soluble α-syn species from MSA, LBD, and Ctrl brains, we measured the viability of differentiated SH-SY5Y cells using the MTT assay. The differentiation of SH-SY5Y cells into neuron-like cells followed an established 10-day protocol using retinoic acid (RA) and brain-derived neurotrophic factor (BDNF) (Suppl. Fig. 7A) (34, 35). An increase of β-III tubulin, MAP2, and synaptophysin levels confirmed the differentiation of the line towards a neuronal phenotype (Suppl. Fig. 7B). We screened the toxicity after a 24-h treatment with serial dilution of α-syn PFF and BH, which demonstrated that 30 ng of PFFs was necessary to observe a significant reduction in viability, whereas no toxicity was observed in PBS brain homogenates except for the LBD-brain derived homogenates when containing 2 ng of α-syn were applied to cells (Suppl. Fig. D,E). When subjecting the cells to either HMW, monomers, or truncated α-syn species, no differences were found between MSA, LBD, and Ctrl cases after 24 h treatment (Suppl. Fig. 7F).

## Discussion

The aim of this study was to characterize brain-derived soluble α-syn species from *post mortem* human brains, comparing differences in seeding and toxicity between α-synucleinopathies that could suggest distinct strain properties.

Previous studies have employed SEC to isolate α-syn species of different sizes from human brain samples, although they have predominantly focused on DLB cases (36, 37) or have employed cross-linking to preserve the conformations of the α-syn species (38–40). It is, however, unclear whether such an intervention introduces a degree of artificiality and alters the biophysical properties of the aggregates (41, 42), making it difficult to draw conclusions about the effects of native α-syn species. To the best of our knowledge, no previous study has directly compared the properties of soluble α-syn species isolated by SEC from both MSA and LBD cases to controls.

A previous study reported no differences in total α-syn levels between individual SEC fractions from cytosolic samples comparing DLB and Ctrl, under non-denaturing and non-cross-linking conditions (36). However, by pooling the levels measured in HMW SEC fractions (∼2000- 440 kDa), the authors reported significantly increased amounts in the DLB group. The amount of HMW α-syn oligomers was reported to be on average 1-1.4% of the total eluted α-syn, indicating that soluble HMW species are present but not abundant (36). Consistent with prior reports, we did not observe any differences in the levels of soluble oligomeric species between α- synucleinopathies and healthy controls across the isolated fractions, indicating that pathological differences are not solely related to quantity. The detection of soluble HMW α-syn species in control brains suggests that oligomeric assemblies are not exclusively associated with disease. α-Syn is increasingly recognized to exist in a dynamic equilibrium between monomeric and self-associated states under physiological conditions, where transient multimerization is thought to contribute to normal synaptic function and protein homeostasis (39, 41, 43).

Several PTMs have been reported in α-syn deposits (22, 23, 44), although their presence and relevance in soluble α-syn species have been understudied. Phosphorylation at Ser129 is the predominant pathological modification of α-syn, with approximately 90% of α-syn within inclusions reported to be phosphorylated in both LBD and MSA (22, 23). Increased pSer129 α-syn was previously reported in MSA RIPA-soluble extracts from the substantia nigra and putamen compared to controls and LBD cases (23). We observed increased phosphorylation in LBD brain-derived HMW oligomers compared to both Ctrl and MSA. Generally, soluble oligomeric α-syn species appeared to be less phosphorylated in MSA than in Ctrl brains, although the difference did not reach statistical significance.

While phosphorylation is considered a secondary PTM, C-terminal truncations have been reported to increase the propensity of α-syn to aggregate and play an active role in the pathogenesis of α-synucleinopathies (45–51). In our SEC fractionation, we collected fractions containing truncated α-syn species and observed increased levels in MSA-derived samples. While there is consensus on increased α-syn truncation in diseases, the observations are generally based on detergent-insoluble preparations, and very few studies have included brain tissue from MSA patients (48, 49). The majority of previous studies assessing α-syn truncation are based on observational immunohistochemical and WB analyses or mass spectrometry (MS) (reviewed in (52)). In the water-soluble brain fraction, C-terminal truncation of α-syn was found to be increased in brains with LB aggregates, although such species were also detected in control brains (46), whereas other studies report no differences (53). Increased α-syn C-terminal truncations were previously reported in MSA RIPA-soluble extracts from the substantia nigra compared to controls and LBD cases (23). The observed differences between our results and those of previous studies may be due to variations in selected brain regions and/or extraction buffers.

The presence of aggregated α-syn has been reported via histopathology and microscopy in LBD and MSA brains (reviewed in (54)). Conventional α-syn immunohistochemistry primarily detects fibrillar inclusions and may underestimate the burden of soluble or diffuse oligomeric pathology. Using α-syn proximity ligation assays (AS-PLA), Roberts et al. demonstrated widespread oligomeric α-syn pathology in PD brain that was largely absent from conventional immunohistochemical analyses and frequently localized outside classical Lewy bodies, suggesting that oligomeric α-syn accumulates before the formation of mature inclusions (55). Similarly, AS-PLA revealed abundant neuronal and oligodendroglial oligomeric α-syn pathology throughout MSA brains, including regions with few detectable glial cytoplasmic inclusions (56). More recently, an aggregate-selective MJFR-14 proximity ligation assay further demonstrated extensive non-inclusion α-syn pathology preceding conventional pSer129-positive Lewy pathology (57). Direct biochemical measurements in MSA and LBD brains have, however, been limited (29, 58). Previous assessments using α-syn conformation-specific antibodies did not report differences between MSA and LBD brains in the substantia nigra (29) nor in the cingulate gyrus, putamen, and amygdala (58). Compared to Ctrl, elevated levels of soluble α-syn oligomers were previously described in DLB (59) and PD brains (60). In our study, the aggregation state of α-syn in cortical PBS brain homogenates discriminated LBD cases from both MSA and healthy controls. The different brain regions analyzed, along with differences in homogenization and normalization processes, complicate direct comparison.

When analyzing specific α-syn sizes, the levels of aggregation in HMW oligomers (>450 kDa) distinguished diseased brains from healthy controls. The observed increase in the LBD group is in line with the finding of a recently published analysis on cortical brain cytosol using MJFR-14- based ELISA, reporting that large HMW oligomers (range of ∼500–1800 kDa) were increased in DLB cases compared to controls (37). Consistent with our findings, total α-syn levels within the oligomeric SEC fractions did not differ between DLB and control brains, whereas the aggregate-specific MJFR-14 ELISA readily distinguished pathological from control samples (37). These findings support the concept that pathological α-syn oligomers possess distinct conformational properties that are selectively recognized by conformation-dependent antibodies despite similar total α-syn levels. While we could measure an increase of HMW oligomers recognized by MJFR- 14 in MSA cases compared to Ctrl cases, their amounts were lower than those of LBDs, and the distribution of aggregated α-syn differed between the α-synucleinopathies, with the maximal α-syn signal appearing in distinct fractions for each disease. The absolute levels (normalized to total protein) of aggregated α-syn, however, did not differ between MSA and LBD in isolated SEC fractions.

The SAA is one of the most prominent strategies for the indirect detection of aggregated α-syn. This assay has shown high sensitivity and, most importantly, can measure functional seeding activities of α-syn species. Historically, SAAs have been highly sensitive but lacked a quantitative capacity, as their kinetic readouts depend on seed conformation and amplification efficiency rather than aggregate abundance (7, 61). While strain differences in SAA have been reported in studies of soluble α-syn species (29, 62), we aimed to understand the specific contribution of differently sized soluble species. The HMW, MMW, and truncated α-syn species derived from both LBD and MSA were capable of initiating aggregation in SAA. Notably, smaller and truncated species displayed pronounced strain-specific differences, with MSA-derived species exhibiting markedly faster and more robust seeding kinetics than their LBD counterparts. While the abundance of aggregated α-syn readily distinguished control from α-synucleinopathy samples, it did not explain the differences in seeding activity observed between LBD and MSA. MSA-derived species isolated via SEC induced a prominent seeded reaction irrespective of their size or aggregation state. Normalized levels of HMW α-syn oligomers from LBD cases, which contained relatively higher levels of aggregated protein, displayed lower seeding potency. The distinct distribution of seeding activity across SEC fractions may reflect underlying conformational differences between LBD- and MSA-derived α-syn species. While our data provide indirect evidence for disease-specific seeding properties, they do not establish structural differences among the soluble assemblies themselves. Cryo-EM studies have demonstrated that mature LBD and MSA fibrils adopt distinct molecular architectures, indicating that the two diseases are associated with different α-syn strains (5, 63). Although it is unknown whether these structural differences extend to soluble oligomeric assemblies, such conformational heterogeneity could influence the size, stability, or templating competence of the minimal seeding species. If analogous structural differences are retained in oligomeric α-syn assemblies, they could contribute to the preferential localization of LBD seeding activity within HMW fractions and the broader distribution of MSA seeding activity observed across SEC fractions. The seeding kinetics were unchanged whether α-syn was normalized or kept at its original concentration within each SEC fraction, highlighting that the conformational and biochemical properties of α-syn species, rather than the aggregation state or abundance alone, determine the observed seeding. Phosphorylation levels also did not correlate with seeding ability.

Seeding activities were also observed in fractions lacking measurable α-syn aggregates, including monomers and truncated species (fractions 17-25), especially for MSA samples. Primary and secondary nucleation reactions could also explain the lack of correlation between aggregate levels and seeding activities in SAA assays. Interestingly, studies of tau have shown that monomeric species can exhibit seeding activity depending on their conformation rather than their molecular size, with distinct inert and seed-competent monomeric states identified (64). Although such a mechanism has not been demonstrated for α-syn, these findings highlight that seeding competence is not necessarily dictated solely by the oligomeric state of a protein.

While a wide size range of α-syn from both LBD and MSA seeded robustly in SAA, only a subset of these species seeded in cell-based models. Soluble brain homogenates from α- synucleinopathies have been analyzed in this and similar models, showing that MSA-derived soluble species induce inclusion formation, whereas soluble α-syn from LBD brains is predominantly inert (8, 65, 66). The MSA-derived HMW oligomers appeared to be the predominant seeding-competent species both in the HEK biosensor line and in murine primary neurons, while species smaller than 450 kDa did not cause seeding or induce aggregation. Similarly, the modest aggregation observed after treatment with LBD-derived samples was limited to the HMW fractions. A previous analysis of DLB-derived HMW oligomers also did not cause aggregation in the HEK biosensor line (36). It is important to consider that the higher seeding ability of MSA-derived α-syn compared to LBD-derived α-syn may be due to strain differences in replication rate, fibril stability, the way the seeds interface with different templates, as well as assay microenvironments (67–69).

Despite their limited seeding activity in cell-based models, soluble α-syn from LBD brains was reported to be toxic to cells. In iDopa cells, 1 ng of α-syn from PBS BH originating from the substantia nigra of LBD patients caused membrane damage and cell death (62). When we applied LBD-derived BH to differentiated SH-SY5Y cells, we observed decreased viability compared to Ctrl, but only with a high amount of α-syn (2 ng). HMW oligomers from DLB cases induced phospholipid membrane permeabilization and caused increased Ca^2+^ influx (36). In the same study, however, SEC-derived HMW oligomers did not affect cell viability in the A53T α-syn HEK293 biosensor line, which was likely attributed to the low α-syn concentration in such samples. In line with previous findings, we did not detect any toxic effect of SEC-isolated α-syn species. Such paradigms, however, can measure acute toxicity while missing the effects of chronic neuronal exposure to α-syn, highlighting the difficulties of replicating year-long diseases in a preclinical setting.

Taken together, our findings suggest that the pathogenic properties of soluble α-syn species are determined primarily by their biochemical, conformational, and functional characteristics rather than by their abundance alone. Although soluble oligomeric α-syn species were present at comparable levels in MSA, LBD, and control brains, disease-associated species displayed distinct biochemical signatures and strain-specific seeding activities that were independent of total α-syn levels. In contrast, control-derived oligomers were not recognized by the aggregate-specific MJFR-14 ELISA and exhibited little or no seeding activity, indicating that oligomerization alone is insufficient to confer pathogenicity. Collectively, these findings support the concept that disease-specific strains are established within the soluble α-syn pool and highlight the importance of targeting disease-associated conformations for the development of more precise biomarkers and therapeutic strategies.

## Materials and methods

### Human tissue samples

Human brain tissues used in this study (three controls, three Lewy body disease, three multiple system atrophy) were provided by the University Health Network Neurodegenerative Disease Collection and were donated under informed consent for research purposes. Autopsy tissues were collected with informed consent from patients or their relatives, in compliance with the approval of the local institutional review board. This study was approved by the University Health Network Research Ethics Board (Nr. 22-5270).

For each brain, one hemisphere was formalin-fixed, dissected into regions, and prepared as formalin-fixed paraffin-embedded tissue sections for neuropathological characterization. The contralateral hemisphere was sliced into ∼1.5 cm thick coronal segments at autopsy, and consecutive slices were formalin-fixed or flash-frozen and stored at −80 °C. This method allows comparison of histopathology in corresponding areas to the frozen samples. Dissection was performed to collect samples from the temporal cortex. Samples were stored in low protein-binding tubes (Eppendorf), flash-frozen, and preserved at −80 °C.

### Protein extraction

Frozen tissues were thawed on wet ice and then immediately homogenized in PBS spiked with protease (Roche, Basel, Switzerland) and phosphatase inhibitors (Thermo Scientific) in a gentle- MACS Octo Dissociator (Miltenyi Biotec) to obtain 10% (w/v) brain homogenates. The homogenate was transferred to a 1.5 mL low protein-binding tube (Eppendorf) and centrifuged at 10,000 g for 10 min at 4 °C, as previously described (29). The supernatant was collected and aliquoted in 1.5 mL low protein-binding tubes (Eppendorf) to avoid excessive freeze–thaw cycles. A bicinchoninic acid protein (BCA) assay (Thermo Scientific) was performed to determine the total protein concentration of all samples. Lysates were stored in low protein-binding tubes (Eppendorf), flash-frozen, and preserved at −80 °C.

### Osmotic shock purification of recombinant α-syn

Full-length, untagged human α-syn was cloned into pET-28a and expressed in E. coli Rosetta 2 (DE3) (Novagen), then purified via osmotic shock and anion exchange as previously described (32, 70, 71). Briefly, expression was induced with IPTG at OD600 >0.6 for ≥3 h. Cells were pelleted (5,000x g, 15 min, 4°C), washed in 1X PBS, and re-pelleted. The pellet was resuspended in osmotic shock buffer (30 mM Tris–HCl pH 7.2, 40% sucrose, 2 mM EDTA; 100 mL per 1000 mL culture), incubated 10 min at room temperature (RT), and centrifuged (9,000 x g, 20 min, 20°C). The resulting pellet was resuspended in ice-cold dH_2_O (40 mL per 1000 mL culture), treated with saturated MgCl_2_ (2.35 g/L; 40 μL per 100 mL suspension), incubated 3 min on ice, and centrifuged (9,000 x g, 30 min, 4°C). The supernatant was collected.

Supernatant was filtered (0.22 μm PES, FroggaBio) and dialyzed overnight at 4°C into 50 mM Tris-HCl pH 8.3 (10K MWCO SnakeSkin™, Thermo Scientific). α-Syn was purified by FPLC on a HiPrep Q HP 16/10 anion exchange column (Cytiva) using a 0–500 mM NaCl linear gradient in 50 mM Tris-HCl pH 8.3. Fractions were assessed by SDS-PAGE followed by Coomassie staining. Pure fractions were pooled, re-dialyzed as above, and further purified on a Mono Q column (GE Healthcare) using the same gradient. After a second SDS-PAGE and Coomassie staining to check purity, pure fractions were dialyzed into dH_2_O. Protein concentration was determined by A280 (NanoDrop; extinction coefficient = 5,960), and aliquots (200 μL) were flash-frozen in liquid nitrogen and stored at -80°C.

Upon thawing and before being used in immunoassays, purified α-syn was filtered through a 100- kDa spin filter (Thermo Scientific) (14,000 x g, 20 min, 4°C) to remove large aggregates; the concentration was measured again, the sample was diluted in TBST (0.05% [*v*/*v*] Tween-20), and was used as a calibrator in the MSD assays.

### Size-exclusion chromatography

Differently sized α-syn species were separated via SEC on a Superdex 200 10/300 GL column (Cytiva) in sterile-filtered PBS, pH 7.4 (Gibco). PBS-solubilized brain extracts (950 µg of total protein) were loaded onto the column and fractionated at a flow rate of 0.5 mL/min using an ÄKTA pure (Cytiva) at 4 °C. A total of 27 fractions (each 0.5 mL) were collected for each sample, starting 6 mL after injection. Fractions were aliquoted and protein concentration was quantified using micro-BCA (Thermo Fisher Scientific) and stored at −80 °C until further use. Dextran Blue (Sigma-Aldrich) was dissolved in phosphate-buffered saline (PBS) at a final concentration of 1 mg/mL and used to determine the void volume of the SEC column. Recombinant monomeric α-syn (rPeptide) was dissolved in ultrapure water at 1 mg/mL and used as a molecular standard to determine the elution volume of monomeric α-syn. Gel Filtration Calibration Kits (Cytiva) were used according to the manufacturer’s instructions to generate a standard curve for molecular weight extrapolation. The partition coefficient (*K*_av_) for each standard was calculated as *K*_av_ = (*V_e_* − *V*_0_)/(*V_t_* − *V*_0_), where *V_e_* is the elution volume, *V*_0_ is the void volume, and *V_t_* is the total column volume. Calibration curves were generated by plotting *K*_av_ against the logarithm of the molecular weight of the standards.

### SDS-PAGE, immunoblotting and Dot-Blot

Gel electrophoresis was performed using 4%–12% Bolt Bis–Tris gels (Thermo Scientific). Proteins were transferred to 0.45-μm nitrocellulose membrane for 60 min at 25 V and crosslinked to the membrane via 0.4% (v/v) paraformaldehyde (PFA) incubation in PBS for 30 min at room temperature (RT), with rocking. The membranes were blocked for 60 min at RT in blocking buffer (5% [w/v] skim milk in 1× PBST (0.05% [v/v] Tween-20) and then incubated overnight at 4°C with primary antibody directed against amino acids 115–122 of the α-synuclein protein (1:1000 dilution, ref: Abcam α/42), diluted in the blocking buffer. The membranes were washed three times with PBST and then incubated, for 60 min at RT, with horseradish peroxidase-conjugated secondary antibodies (172-1011, Bio-Rad) diluted 1:2000 in the blocking buffer. Following another three washes with PBST, immunoblots were developed using the SuperSignal^TM^ West Pico PLUS chemiluminescence substrate (Thermo Scientific) and imaged using X-ray film.

For Dot-Blot analyses, 1 µL from each SEC fraction was pipetted on a nitrocellulose membrane and allowed to air-dry for 60 min. The membranes were then fixed with 0.4% PFA and immunoblotted as described above. For the detection of aggregated α-syn, the conformation-specific antibody MJFR-14 (Abcam) was used at a 1:2000 dilution. pSer129-syn was probed with D1R1R antibody (Cell Signaling) at a dilution of 1:2000.

### Total α-syn ELISA and Electrochemiluminescent MSD assays

Total human α-syn ELISA kits (Anaspec) were used following the manufacturer’s protocol.

The MSD technology platform (Mesoscale Discovery) was used to develop electrochemiluminescent assays for pSer129 α-syn and aggregated α-syn. . MJFR-13 (Abcam, ab168381), MJFR-14 (Abcam, ab209538), and MJFR1 (Abcam, ab138501) were labeled with biotin and SULFO-Tag using the EZ-Link Micro Sulfo-NHS-LC Biotinylation kit (Thermo Scientific) and MSD GOLD SULFO-TAG NHS-Ester Conjugation Pack (Mesoscale Discovery), following the manufacturer’s instructions.

The development of the pSer129 α-syn immunoassay was performed with minor modifications from previously published protocols (72, 73). Semisynthetic pSer129 α-syn (Proteos Inc.) was used as a standard and prepared as previously described (74). MSD Gold 96-well Small Spot streptavidin plates were coated with 1µg/mL (diluted in Diluent 100, Mesoscale Discovery) of the biotinylated capture antibody (MJFR-13), sealed with an adhesive foil, and incubated for 1 h at RT with shaking (750 rpm). After three washes with 150 µL of TBS-T (0.05% v/v Tween-20), 25 µL of a 1µg/mL dilution of the detection antibody (Sulfo-TAG MJFR1) and 25 µL of each standard and sample were added to the respective wells. Detection antibody, standards, and samples were diluted in TBS containing 0.1% (v/v) Tween-20 (TBS-T) and 1% BSA. The plate was incubated for 2 h under shaking conditions (750 rpm) and protected from light before being washed three times. 150 µL of Gold Read Buffer B (Mesoscale Discovery) was added to each well, and the plate was read on an MSD Quickplex Q60 reader and analyzed with the MSD Methodical Mind Analysis Software.

For the development of the aggregated α-syn-specific assay, protocols were adapted from previous publications (37, 75). Recombinant α-syn PFF (rPeptide), were diluted to 20 μg/mL in TBS containing 1% BSA, sonicated for 5 min at 70V with a QSonica sonicator, aliquoted, and stored at -80 °C as a stock standard solution. The dilution of the antibodies (biotinylated and sulfo-TAG MJFR-14), standards, and sample in this assay was performed in Diluent 49 (Mesoscale Discovery). The assay procedure is the same as for the pSer129 α-syn assay.

Further details on the development and validation of the immunoassays are described in the supplementary information (Table S2, Suppl. Fig.2)

### α-Syn seeding amplification assay (SAA)

The SAA reactions were performed in 384-well plates with a clear bottom (Corning) with minor modifications from previously described protocols (29, 62). Briefly, full-length human recombinant α-syn with the K23Q mutation (Impact Biologicals) was thawed from − 80°C storage, reconstituted in HPLC-grade water (Sigma), and filtered through a 100-kDa spin filter (Thermo Scientific) in 500 µl increments. Ten µl of the biological sample (5 µg of total protein from the PBS BH, 1 ng α- syn from SEC fractions) was added to the wells containing 20 µl of the reaction buffer (0.1 M phosphate buffer, pH 8, 0.425 M Na_3_Citrate), 10 µl of 50 µM ThT and 10 µl of 0.5 mg/ml of monomeric recombinant K23Q α-syn. In the analysis of SEC fractions, the reactions were spiked with PBS BH from triple-syn KO mice (kindly provided by Prof. Sreeganga Chandra, Yale University) at an in-assay concentration of 40 µg/mL. The plate was sealed and incubated at 42 °C in a BMG FLUOstar Omega plate reader with cycles of 1 min shaking (400 rpm double orbital) and 1 min rest. ThT fluorescence measurements (450 ± 10 nm excitation and 480 ± 10 nm emission, bottom read) were taken every 15 min for 120 h. Each sample was tested in triplicate and was considered positive if 2 out of 3 replicates generated the expected sigmoidal-shaped curve for a positive seeding reaction. SAA fluorescence curves were fitted using a four-parameter logistic (sigmoidal) model in GraphPad Prism.

### HEK293 α-syn(A53T)-YFP biosensor line

HEK293 cell line stably expressing A53T-mutant human α-syn tagged at its C-terminus with yellow fluorescent protein (YFP) was cultured at 37°C and 5% CO_2_ in Dulbecco’s modified Eagle’s medium (DMEM) supplemented with 10% fetal bovine serum (FBS), 100 U/ml penicillin, 100 μg/ml streptomycin, 2 mM l-glutamine, 1 mM sodium pyruvate. For the protein inclusion formation assays, cells were plated overnight on a black 96-well, clear-bottom plate (Thermo Scientific) at a density of 30,000 cells/well, previously coated with 0.05 mg/mL Poly-D Lysine (Gibco). Treatment was performed as previously described (8, 36). Samples (50 ng of α-syn PFF, 10 µg BH, 25 µL SEC fractions) were diluted in Opti-MEM with 2% (v/v) Lipofectamine 2000 and incubated at RT for 1 h. Culture medium was removed, and the cells were incubated with the sample-Lipofectamine mixtures for 4 h. After completing the treatment, the mixtures were removed and replaced with fresh culture medium without antibiotics. The cells were then cultured for 96 h before being fixed with 4% PFA and counter-stained with DAPI. Images were taken at 40x magnification using a Nikon Confocal (NIS-Elements software). The number of aggregates was counted after applying a fixed threshold on Fiji and expressed as a percentage of DAPI-positive cells.

### Primary Hippocampal Culture

Primary neuronal cultures were prepared from E16 CD1 mouse brains (Charles River, Quebec, Canada). All procedures were performed according to the Canadian Council on Animal Care and the Animals for Research Act of Ontario. Brains of E16 mouse pups were retrieved, and the hippocampal regions were dissected and collected in HBSS solution. The collected hippocampal portions were pooled into a tube and treated with 0.035% trypsin in a water bath at 37°C. After 30 min, 5 mL of 0.0001% DNAase 1 solution was added. After removing the DNase solution, 500 µl of trituration buffer was added, and the cells were triturated with polished Pasteur pipettes 10 times. After adding 5 mL Neurobasal medium, the preparation was filtered through a 0.65 µm filter and counted and plated at 100,000 cells/well in 24-well plates.

Cells were maintained in Neurobasal medium supplemented with B27, 1 mM glutamate, and 50 U/ml penicillin/streptomycin at 37°C in a 5% CO2 incubator. After 7 days of plating, cells were treated with pooled SEC fractions, normalized to 1 ng of total α-syn, in 200 µL of media; after 24 h, 200 µL more conditioned media was added. Cells were fixed 14 days after treatment.

### Immunocytochemistry and quantification

Neurons were washed 2-3x with D-PBS, then fixed with 4% paraformaldehyde (PFA) and 4% sucrose in PBS for 15 min. The PFA was then washed off three times with PBS, and cells were permeabilized with 0.2% Triton X-100 for 5 min. Subsequently, cells were blocked for 1 hour in 5% goat serum followed by a 1-hour incubation with primary antibodies diluted in blocking buffer. The following antibodies were used: anti-pSer129 α-syn antibody (monoclonal, EP1536Y, 1:500, Abcam), anti-MAP2 (monoclonal, HM-2, 1:1000, Sigma). A 1-hour incubation with species-specific secondary antibodies coupled to Alexa Fluor 488, 594 (1:500, Abcam) followed. Coverslips were then immediately incubated with 4’, 6-diamidino-2-phenylindole (DAPI) (D9542, 1:10,000, Sigma-Aldrich) for 5 min followed by three washes with PBS. Coverslips were then mounted onto Fisherbrand Superfrost Plus microscope slides (Thermo Fisher Scientific) using DABCO mounting media (Sigma).

The following method was used to quantify the pSer129 α-syn pathology: 5 randomly selected visual fields from each condition were taken on the confocal microscope at 40x, zoom 1. In ImageJ, images were converted to binary, and then fixed thresholds were set for p-α-syn and MAP2 channels within each experiment; p-α-syn was quantified using the Analyze Particles function, with pixel size set at 0.01-infinity, to remove background. The area output of p-α-syn staining was then normalized to the area of MAP2, and ratios were graphed.

### SH-SY5Y – differentiation, treatment, and toxicity assay

SH-SY5Y human neuroblastoma cells (ATCC CRL-2266) were cultured in DMEM/F12 with 10% FBS, 100 U/ml penicillin, 100 µg/ml streptomycin, and 2 mM L-glutamine at 37°C and 5% CO_2_. The cells were differentiated towards a neuron-like phenotype following an established 10-day protocol using retinoic acid (RA, Thermo Scientific) and BDNF (Thermo Scientific) (34, 35). 24h prior to the beginning of the differentiation procedure, cells were plated in a 96-well plate at 2.0 x 10^4^ cells/well in complete growth medium. The growth medium was then replaced with differentiation medium containing 10 µM RA in DMEM/F12 with reduced serum (1%), which was refreshed after 72h. On day 7, the differentiation medium was replaced with maturation medium containing 50 ng/mL BDNF in DMEM/F12 with reduced serum (1%). On day 9, the maturation medium was refreshed, and treatment with protein samples started. After 24 or 48h, the sample-containing media was removed, and the CyQUANT™ MTT Cell Viability Assay (Thermo Scientific) was performed following the manufacturer’s instructions. Cells treated with 0.1% Triton X-100 served as the positive control for complete loss of cell viability in the MTT assay. Experiments were performed in triplicate.

## Acknowledgments

Prof. Sreeganga Chandra, Yale University, kindly provided snap-frozen brain hemispheres from triple α-syn KO mice. The authors thank M. Diamond (UT Southwestern) for providing the HEK293 cells expressing YFP-tagged α-syn(A53T). We would also like to acknowledge the Edmond J Safra Program in Parkinson Disease for providing funding for the maintenance of the University Health Network Neurodegenerative Disease Collection.

## Supplementary information

**Table S1.** Neuropathological diagnosis of the subjects included in this study. Summary of the neuropathological and clinical characteristics of the subjects analyzed. Alzheimer’s disease neuropathological change is reported using the NIA-AA “ABC” score (A, Thal phase of Aβ deposition; B, Braak neurofibrillary tangle stage; C, CERAD neuritic plaque score). Abbreviations: Aβ, amyloid-β; CERAD, Consortium to Establish a Registry for Alzheimer’s Disease; F, female; LATE-NC, limbic-predominant age-related TDP-43 encephalopathy neuropathological change; M, male; LBD, Lewy body disease; MSA, multiple system atrophy; NIA-AA, National Institute on Aging–Alzheimer’s Association.

| Case | Sex | Age | Disease | NIA-AA | Cerebral Amyloid Angiopathy | Lewy Body Braak Stage | LATE-NC Stage |
| --- | --- | --- | --- | --- | --- | --- | --- |
| MSA 1 | F | 68 | MSA | A1B1C0 | Aβ-positive Type 2 | No | No |
| MSA 2 | M | 72 | MSA | A1B0C0 | No | No | No |
| MSA 3 | M | 64 | MSA | A0B1C0 | No | No | No |
| LBD 1 | M | 73 | LBD | A2B3C2 | Aβ-positive Type 2 | Stage 5 | Stage 2 |
| LBD 2 | M | 78 | LBD | A3B2C2 | Aβ-positive Type 2 | Stage 5 | Stage 2 |
| LBD 3 | M | 76 | LBD | A2B1C1 | No | Stage 4 | No |
| Ctrl 1 | F | 48 | Control | A0B0C0 | No | No | No |
| Ctrl 2 | M | 74 | Control | A0B0C0 | No | No | No |
| Ctrl 3 | F | 69 | Control | A1B1C1 | No | No | No |

**Supplementary Fig. 1.**
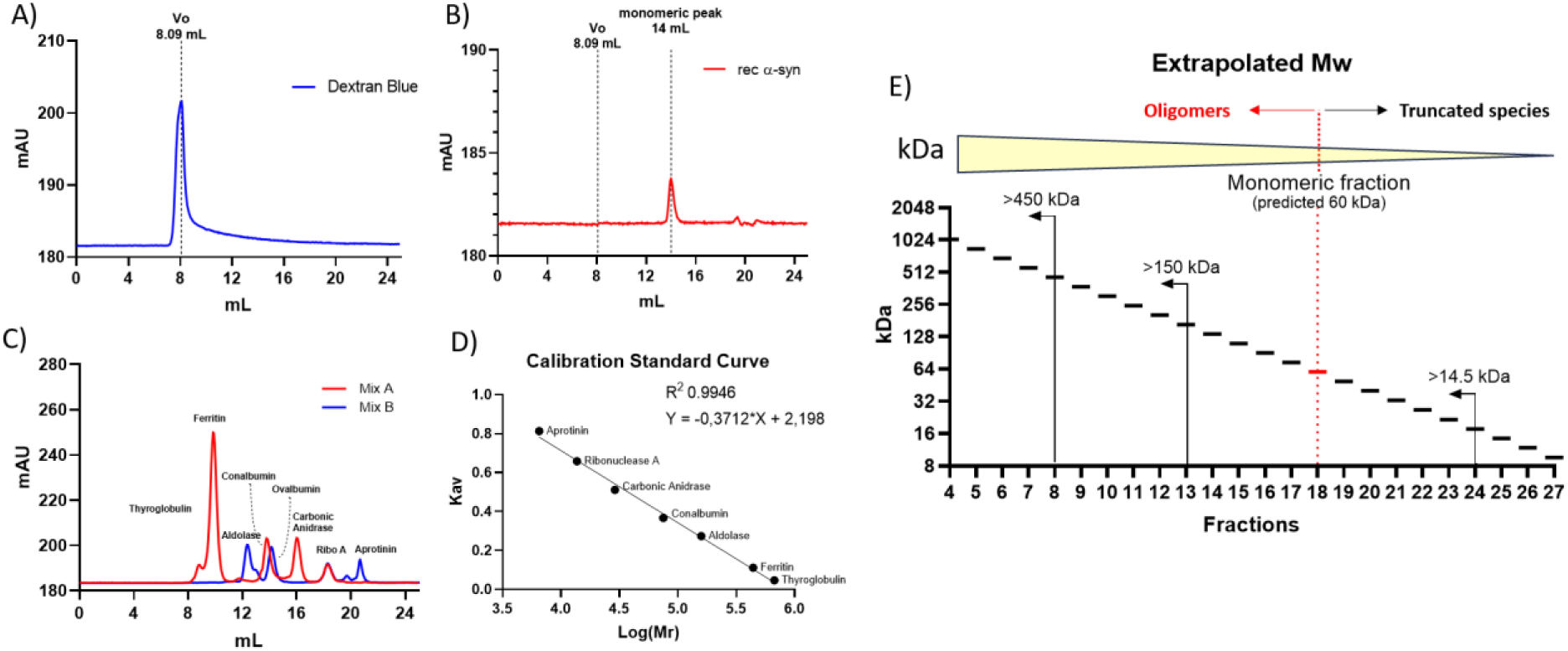
A) Void volume (Vo) determination. A 1 mg/mL solution of Dextran Blue 2000 was injected to determine the column void volume using the earliest-eluting high-molecular-weight species. B) Recombinant α-synuclein standard. Purified WT rec α-synuclein was run to define the elution position of the monomeric species under chromatographic conditions. C) Calibration standards. A mixture of Cytiva HMW and LMW gel filtration calibrators (Mix A and Mix B) was injected to generate reference elution times for proteins spanning a broad molecular-weight range. D) Calibration curve. A standard curve was generated by plotting the log₁₀(MW) of each calibrator protein against its corresponding Kav, yielding a linear relationship used for molecular-weight estimation. E) Molecular-weight extrapolation of sample fractions. The calibration curve was used to extrapolate the Mw range for each collected fraction, thereby assigning approximate molecular weights and classifying species as monomers, oligomers, or truncated/low-MW based on their elution positions.

### Development of pSer129 α-syn and aggregated α-syn electrochemiluminescence immunoassays

Aliquots of mAb MJFR13 (specific towards pSer129 α-syn) and mAb MJFR14 (specific towards aggregated α-syn) were conjugated with biotin to serve as capture antibodies. Aliquots of mAb MJFR1 (serving as total syn-mAb) and MJFR14 (specific towards aggregated α-syn) were conjugated with Sulfo-Tag to serve as detection antibodies. For the development of the pSer129- syn specific assay, 96-well small spot streptavidin plates were coated with biotinylated mAb MJFR13 and paired with Sulfo-TAG mAb MJFR1, both used at a concentration of 1 µg/mL. Semisynthetic pSer129-syn (Proteos Inc.) was used to generate a standard curve. For the development of an aggregates α-syn specific assay, the MJFR14 antibody (1 µg/mL) was used both as capture and detection antibody. PFF from rPeptide were used to generate standard curves.

### Lower Limit of Detection, Lower Limit of Quantification, and Upper Limit of Quantification

The measurements in each assay run were derived from fitting a four-parameter logistic standard curve using MSD Methodical Mind Analysis Software. The raw signals were obtained from seven fourfold serial dilutions of the recombinant α-syn calibrators plus a zero calibrator (blank). An initial extended fourfold calibrator dilution series, including ten fourfold serial dilutions starting at 300 ng/ml, was analyzed to determine the upper limit of quantification (ULOQ) and the appropriate standard curve range for each assay. The ULOQ was measured in a single assay and is defined as the highest calibrator concentration that showed < 20% coefficient of variation (CV) between two technical replicates (regarding both signal and calculated concentration) and 85–115% recovery (i.e., the calculated concentration had to be in the range of 85–115% of the theoretical calibrator concentration). Additionally, the LLOD was calculated for each assay plate as the lowest analyte concentration that generates a signal 2.5 SDs above the lowest standard/blank, with the option “use minimum error estimates” activated. The lower limit of quantification (LLOQ) was defined as the lowest analyte concentration producing a signal 10 SDs above the lowest standard/blank. ULOQs and mean LLODs and LLOQs, calculated from n = 3-4 assay runs, are summarized in **Table S2**. The starting concentration of the fourfold dilution pSer129 α- synstandard curve was chosen at 18750 pg/ml, while 120000 pg/ml was the starting concentration of the PFF calibrator used in the aggregated α-syn specific assays.

**Table S2:** Average LLODs, LLOQs and ULOQ of the α-syn-specific immunoassays.

| α-Syn assay | Capture mAb-Biotin | Detection mAb-sulfo-Tag | LLOD | | LLOQ | | ULOD |
| --- | --- | --- | --- | --- | --- | --- | --- |
|  |  |  | pg/mL | SD | pg/mL | SD | pg/mL |
| pSer129 α-syn | MJFR-13 | MJFR-1 | 2.669 | 1.50 | 27.877 | 8.00 | 300000 |
| Aggregated α-syn | MJFR-14 | MJFR-14 | 26.225 | 5.07 | 105.572 | 39.02 | 300000 |

### Cross-reactivity with unphosphorylated and monomeric α-syn

The MJFR-13 and MJFR-14 antibody specificity towards phosphorylated aggregated was assessed in a dot blot analysis prior to the development of the MSD assays. Serial dilution of pSer129 α-syn, WT rec α-syn monomers, and α-syn PFF used as standards in the assays, as well as SEC fractions containing aggregated and monomeric α-syn, were loaded onto nitrocellulose membranes and probed with the primary mAb as described in the Materials and Methods section. As shown in Suppl. Fig 2A,B, the MJFR13 antibody specifically recognized pSer129 α-syn standard, while the MJFR1 mAb bound to both phosphorylated and unphosphorylated α-syn. MJFR14 recognized species in aggregate-containing SEC fractions as well as PFF, but did not bind to monomers in fraction 18 Suppl. Fig 2D. The specificity was also confirmed in the MSD assays. A standard curve of unphosphorylated rec α-syn was measured in the pSer129 assay, showing that until 1171.9 pg/mL, unphosphorylated α-syn was below the detection limit, while a 0.2% cross-reactivity was measured at concentrations of 4687.5 and 18750 pg/mL (Suppl. Fig 2B,C). In pilot measurements with the aggregated-α-syn assay, no α-syn could be detected in fractions containing monomeric α-syn, while species were recognized in fraction 4, containing aggregates (Suppl. Fig 2E).

**Supplementary Fig. 2.**
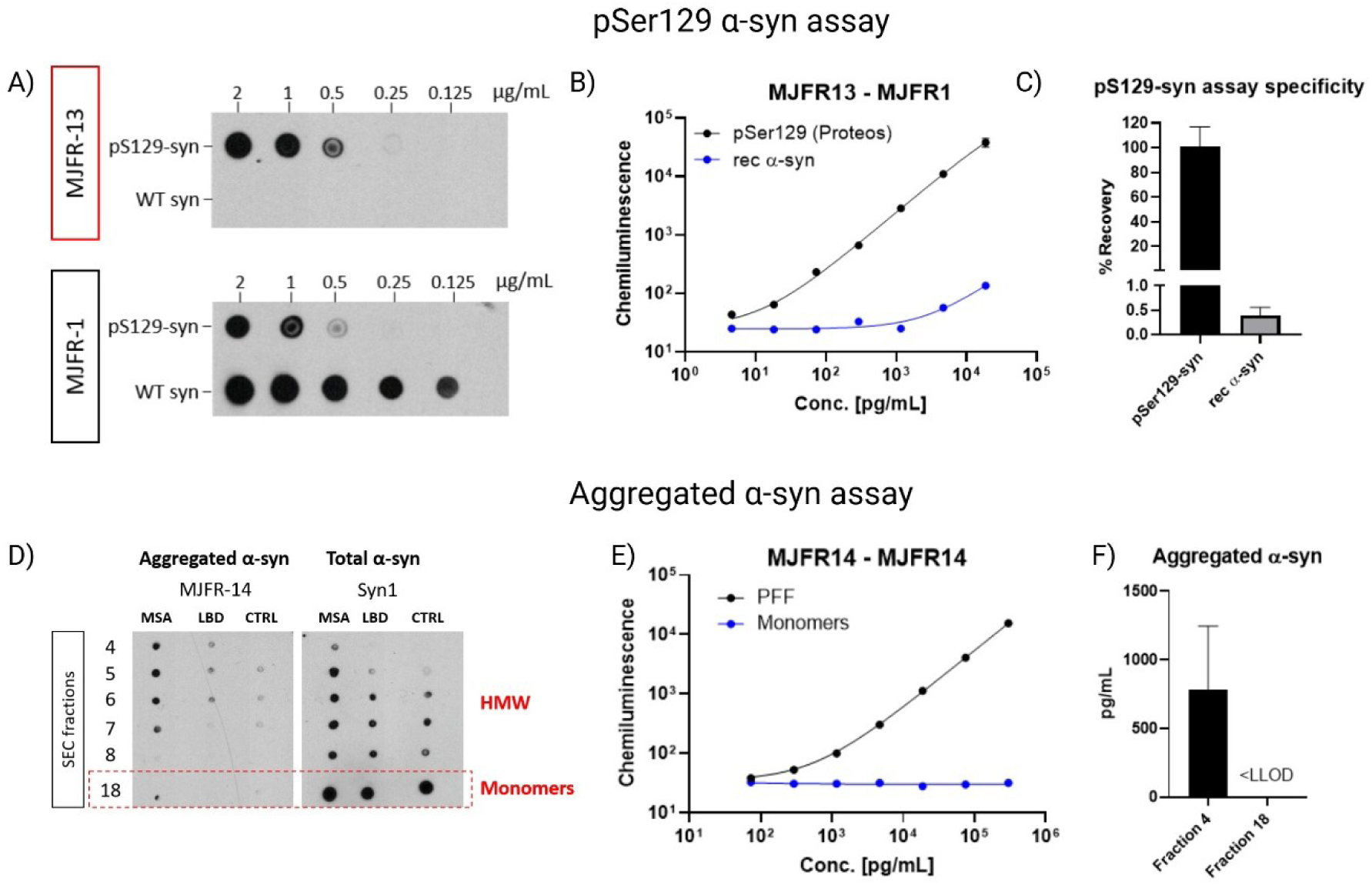
A) Dot blot specificity of MJFR-13 and MJFR-1. Serial dilutions of phosphorylated and unphosphorylated α-synuclein were prepared to assess antibody selectivity for pSer129 versus total α-syn. B) pSer129 MSD immunoassay specificity. Calibration curves were generated using semisynthetic pSer129-α-synuclein and unphosphorylated recombinant α-synuclein. The mAb combination recognized the expected concentration-dependent signal and confirmed selective recognition of the phosphorylated epitope. C) Recovery of unphosphorylated α-syn. The percent recovery of non-phosphorylated α-synuclein in the assay indicates minimal cross-reactivity and quantifies background binding. D) Dot-blot validation of MJFR-14. Aggregated α-synuclein species contained in fractions 4-8, but not monomers, were recognized by the conformation-specific antibody MJFR-14. E) Aggregated α-synuclein MSD immunoassay specificity. Calibration curves were generated using rec PFF and osmotic shock WT rec α-syn monomers. The mAb combination recognized the expected concentration-dependent signal and confirmed selective recognition of aggregated α-syn. F) The aggregated α-syn assay detects aggregated α-synuclein in fraction 4 (n=3) and shows no detectable binding to monomeric α-synuclein in fraction 18 (n=3).

**Supplementary Fig. 3.**
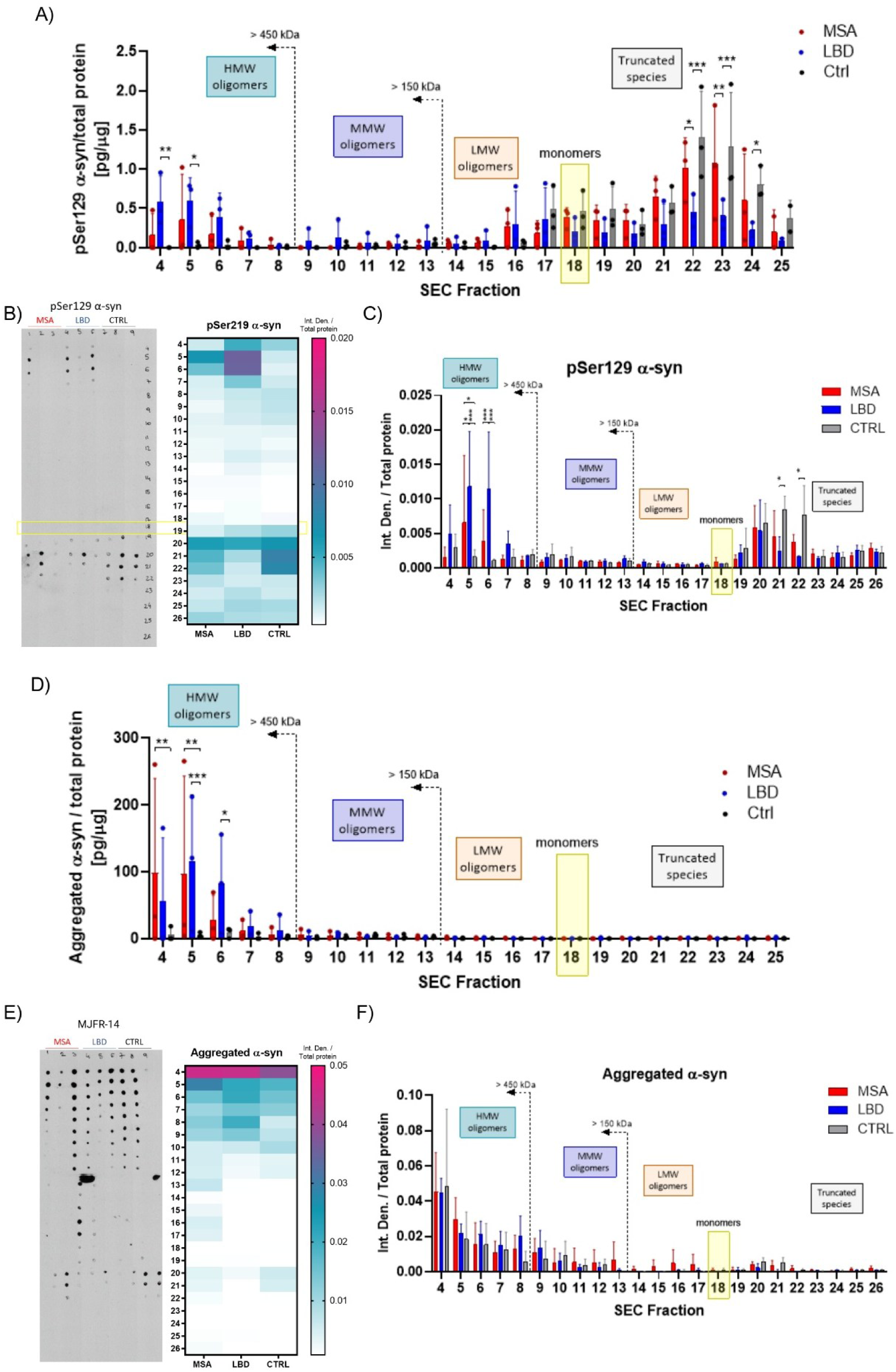
A) Levels of pSer129 α-syn in each SEC fraction, measured with MSD immunoassay, normalized to the amount of total protein. B) Dot-blot detection of pSer129-α-syn across SEC fractions. A heatmap of spot intensity illustrates the relative abundance of pSer129-α-syn across fractions. C) Densitometric analysis of dot-blot signals normalized to total protein. D) Levels of aggregated α-syn in each SEC fraction, measured with MSD immunoassay, normalized to the amount of total protein. E) Dot-blot detection of aggregated α-syn across SEC fractions, visualized as a heatmap of signal intensity across fractions. F) Densitometric analysis of aggregated-syn dot-blot signals normalized to total protein.

**Supplementary Fig. 4.**
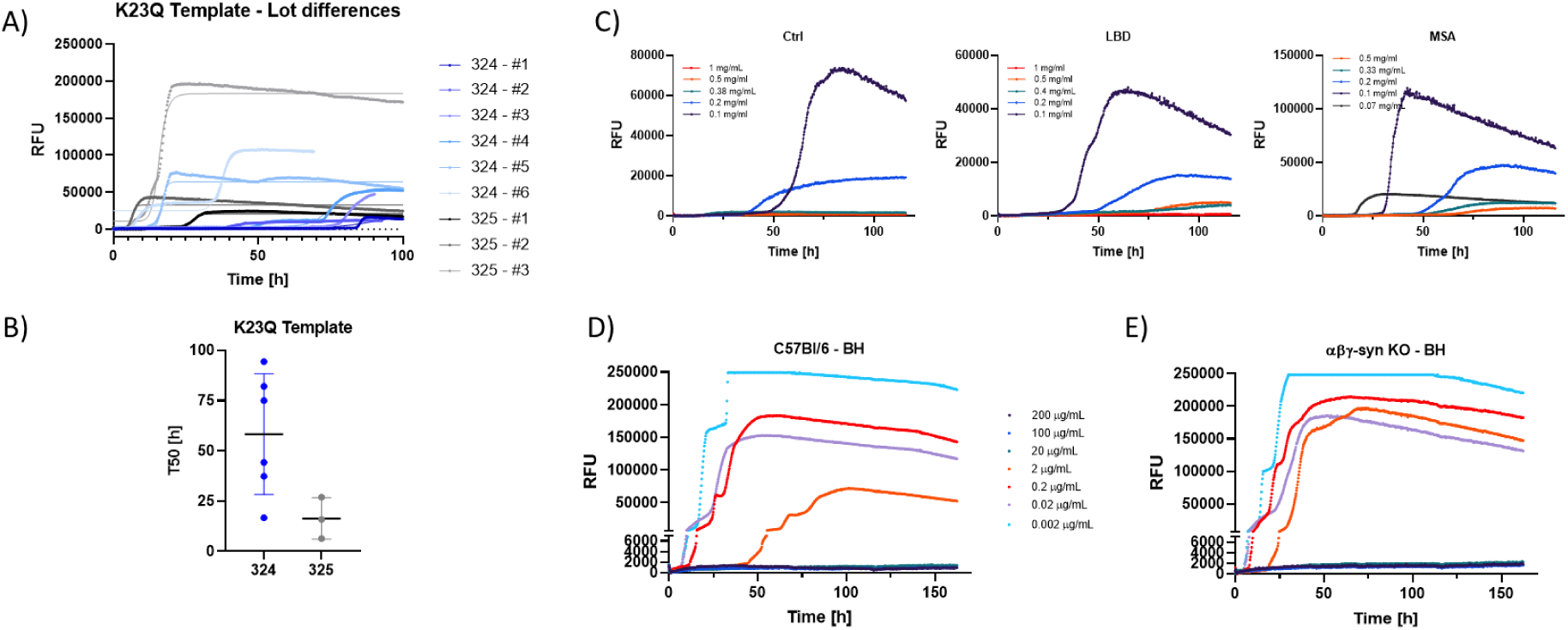
A) Spontaneous aggregation kinetics of α-synuclein SAA using the K23Q mutant template. Two lots of template (324 and 325) were analyzed in independent SAAs runs (#1-6). B) T50 values demonstrating intra- and inter-lot variability in template spontaneous aggregation. C) SAA response to serial dilutions of human brain homogenate (BH) from Ctrl, LBD, and MSA cases. D) SAA with serial dilutions of WT mouse (C57BL/6) brain homogenate. E) SAA with serial dilutions of triple-synuclein knockout (α-, β-, γ-syn KO) brain homogenate.

**Supplementary Fig. 5.**
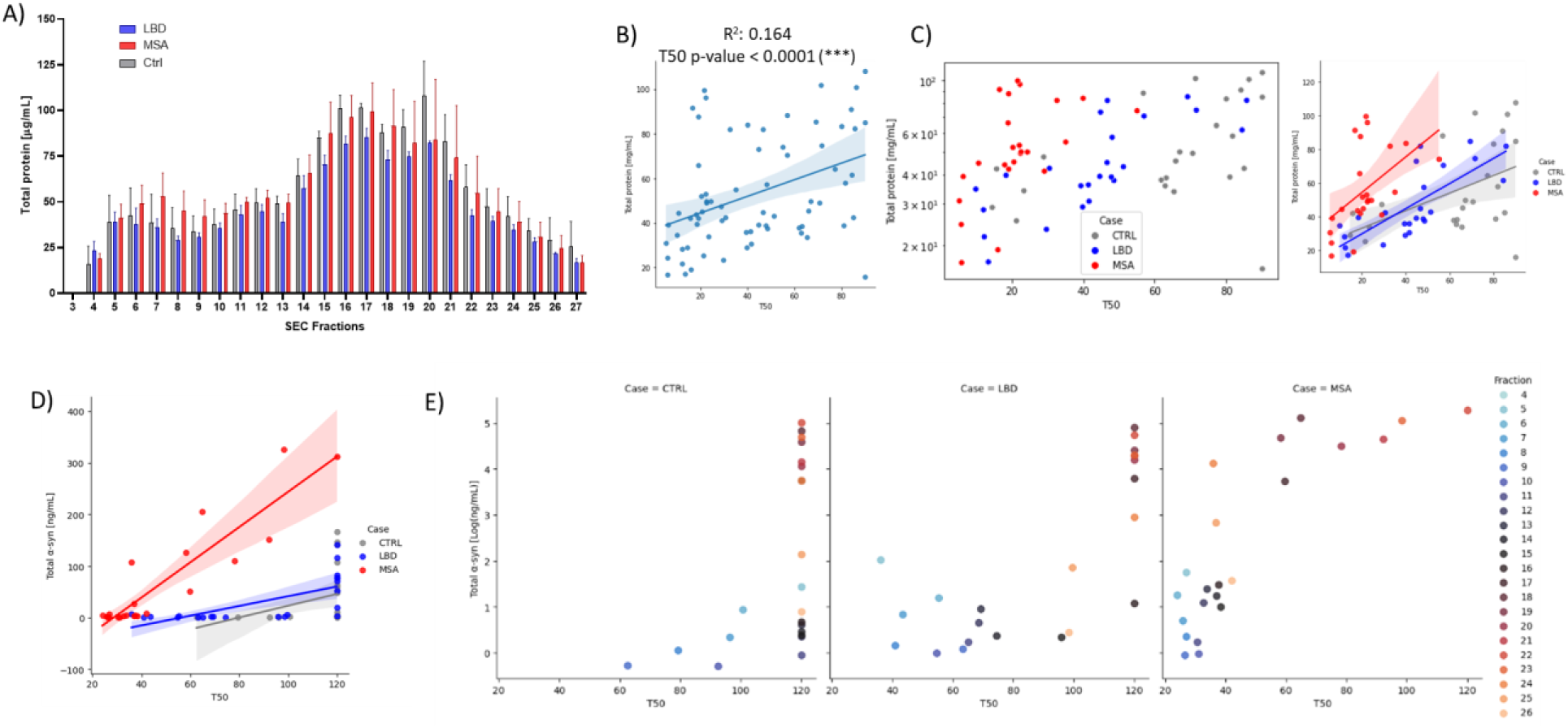
A) Total protein concentration across SEC fractions measured by µBCA. B) Scatterplot showing the relationship between SAA T50 values and total protein levels across all SEC fractions in the absence of triple-syn KO BH top-up. C) Case-stratified correlation plots illustrating how the T50–protein relationship varies across diagnostic groups. D) Scatterplot showing the association between SAA T50 values and total α-syn levels measured in each SEC fraction. SAAs were performed with undiluted SEC fractions and with the addition of triple-syn KO BH to reach 40 µg/mL total protein concentration. E) Case-grouped relationship between α-syn concentration and T50 across SEC fractions from D.

**Supplementary Fig. 6.**
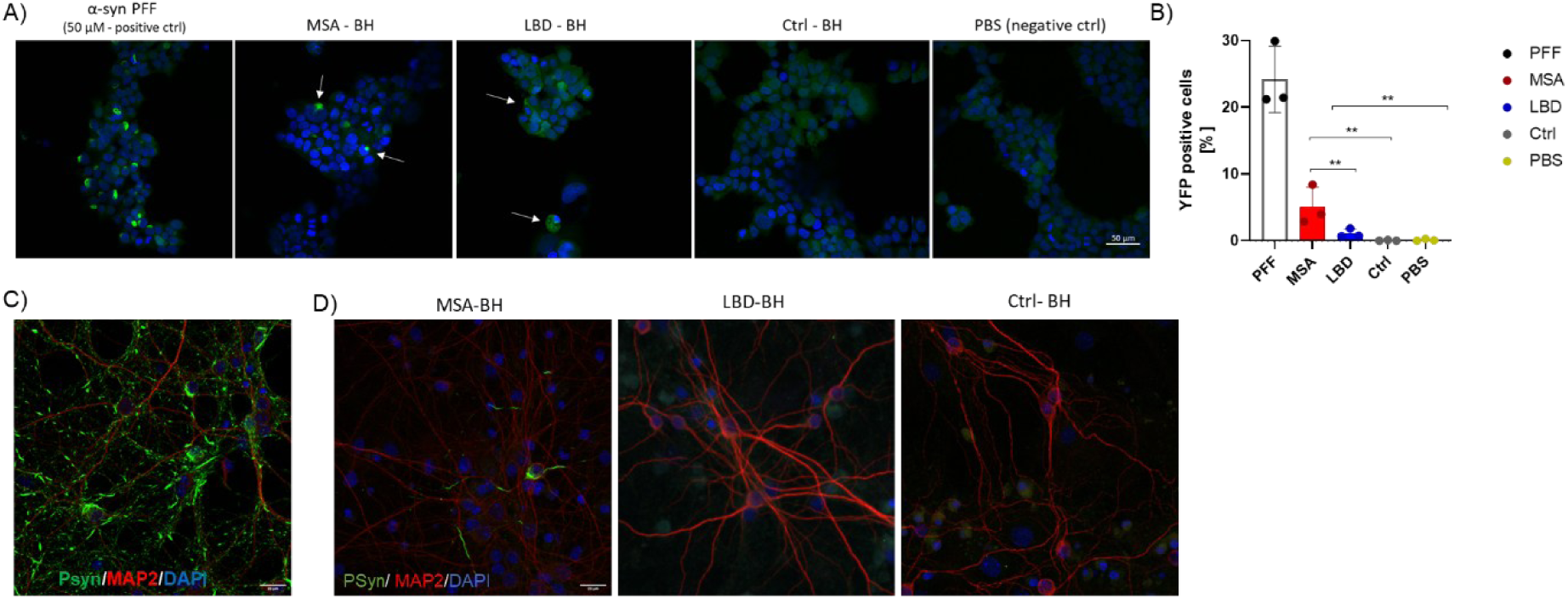
A) Representative images of HEK293T α-syn biosensor cells following exposure to PFFs, brain homogenate, or PBS, with arrowheads indicating YFP-positive aggregates. B) Quantification of seeding activity in biosensor cells. Percentage of YFP aggregate-positive cells across conditions demonstrates strong seeding in PFF-treated and MSA-treated wells, minimal signal in LBD, and baseline levels in CTRL and PBS controls. C) pSer129-α-syn (Psyn, green) accumulation in WT primary neurons exposed to α-syn PFFs. D) Representative images from murine primary neurons treated with Ctrl, LBD, and MSA brain homogenates, illustrating disease-dependent differences in pSer129-positive inclusion formation. Microtubule-associated protein (MAP2, red) marks neuronal morphology; DAPI (blue) counterstains nuclei.

**Supplementary Fig. 7.**
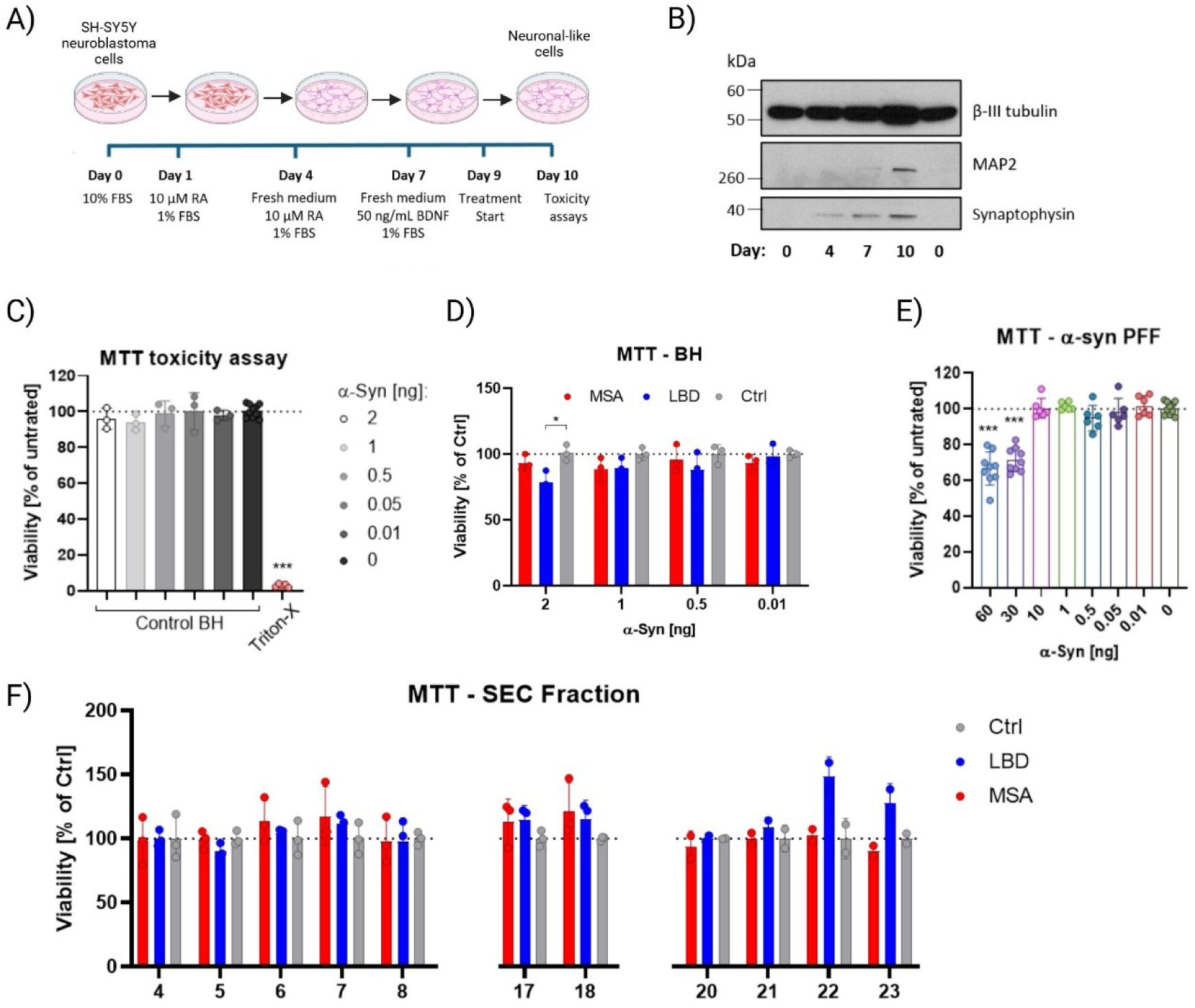
A) Schematic of the differentiation protocol using RA and BDNF, illustrating the timeline of media changes and maturation steps leading to a neuron-like phenotype (Adapted from (35)). B) Western blot validation of neuronal differentiation. Representative immunoblots showing progressive increases in βIII-tubulin, MAP2, and synaptophysin expression across the differentiation timeline. C) MTT viability assay with control brain homogenate. SH-SY5Y viability remains unchanged following exposure to CTRL brain homogenate, whereas Triton X-100 treatment induces marked cell death. D) MTT assay with serial dilutions of brain homogenate normalized by α-syn content. E) MTT viability assay with α-syn PFFs, demonstrating concentration-dependent toxicity. F) MTT assay with SEC-derived α-syn species, expressed as % of Ctrl-treated samples.

